# The role of *Staphylococcus agnetis* and *Staphylococcus hyicus* in the pathogenesis of buffalo fly skin lesions in cattle

**DOI:** 10.1101/2022.03.11.483979

**Authors:** Muhammad Noman Naseem, Conny Turni, Rosalind Gilbert, Ali Raza, Rachel Allavena, Michael McGowan, Constantin Constantinoiu, Chian Teng Ong, Ala E. Tabor, Peter James

## Abstract

Buffalo flies (*Haematobia irritans exigua*) are hematophagous ectoparasites of cattle causing production and welfare impacts in northern Australian herds. Skin lesions associated with buffalo fly infestation and *Stephanofilaria* nematode infection are manifested as focal dermatitis or ulcerated areas most commonly on the medial canthus of the eye, along the lateral and ventral neck and on the abdomen of cattle. For closely related horn flies (*Haematobia irritans irritans*), *Staphylococcus aureus* have been suggested as a contributing factor in the development of lesions. To investigate the potential role of bacterial infection in the pathogenesis of buffalo fly lesions, swabs were taken from lesions and normal skin, and bacteria were also isolated from surface washings of buffalo flies and surface-sterilised homogenized flies. Bacterial identification was conducted by MALDI-TOF, strain typing by rep-PCR and DNA sequencing to determine species similarity and virulence factors. Of 49 bacterial isolates collected from lesions, 37 were identified as *Staphylococcus agnetis* and 12 as *Staphylococcus hyicus*, whereas from normal skin four isolates were *S. hyicus* and one was *Staphylococcus sciuri*. Of the *Staphylococcus* isolates isolated from buffalo flies, five were identified as *S. agnetis* and three as *S. hyicus*. Fifty percent of the buffalo fly isolates had rep-PCR genotypic patterns identical to the lesion isolates. Genome sequencing of 16 *S. agnetis* and four *S. hyicus* isolates revealed closely similar virulence factor profiles, with all isolates possessing exfoliative toxin A and C genes. The findings from this study suggest the involvement of *S. agnetis* and *S. hyicus* in buffalo fly lesion pathogenesis. This should be taken into account in the development of effective treatment and control strategies for lesions.

## INTRODUCTION

Buffalo flies (BFs) (*Haematobia irritans exigua*) are hematophagous ectoparasites, closely related to horn flies (HFs) (*Haematobia irritans irritans*), which feed mainly on cattle and buffaloes (1, 2, 3). Buffalo flies occur in the tropical and subtropical parts of Australia, Asia, and other parts of Oceania, while HFs are prevalent in South and North America and Europe (4). In Australia, cattle skin lesions associated with buffalo fly (BF) feeding are termed BF lesions. These lesions can range from raised, dry, alopecic, hyperkeratotic or scab encrusted areas to severe open suppurating wounds occurring mainly near the medial canthus of the eye, neck, and ventral midline (5). Although these lesions are associated with BF feeding, Sutherst et al., reported a low correlation between BF counts and lesion development (6).

An unnamed species of *Stephanofilaria* nematode has been implicated in the development of these lesions (5), but nematodes were detected in only 40% of skin lesions (7). Naseem et al., suggested that *Stephanofilaria* sp. infection might not be essential for BF lesion development as their study found only 10.83% of lesions infected with *Stephanofilaria* sp., with no nematodes found in either lesions or BFs in some regions of Australia despite the frequent occurrence of lesions (Naseem et al., submitted for publication). Horn flies are reported as vectors for *S. stilesi* nematodes, which have also been implicated in the development of skin lesions in cattle in north and south America (1). However, hypersensitivity to horn fly (HF) feeding and the involvement of *S. aureus* have also been suggested as contributing causes in the development of these lesions (8, 9). These findings suggest that other factors might be involved in the development of BF lesions.

In the USA, HF have been identified as vectors of *Staphylococcus aureus* bacteria, which have been isolated from lesions on the teats and udders of dairy cattle (9). Nickerson et al., showed that dairy farms using HF control presented lower rates of *S. aureus* udder infection compared to herds without control (10) and later Gillespie et al., confirmed that *S. aureus* isolates from HF had identical DNA fingerprints to 95% of *S. aureus* isolates from udder skin lesions (11). In addition, *S. aureus, S. saprophyticus, S. hyicus* and *S. sciuri* have been identified in the microbiome of HF (12).

*Staphylococcus hyicus* has also been isolated from fresh, encrusted, dry and old healing skin lesions on the back, shoulder, and root of the tail of cattle, and experimental inoculation with *S. hyicus* produced lesions with a similar clinical appearance (13). Hazarika et al., also isolated *S. hyicus* from skin lesions around the eye, forehead, neck, shoulder, hump, and trunk of cattle, and reproduced skin lesions in rabbit skin by experimental inoculation with isolated *S. hyicus* (14). *Staphylococcus hyicus* has also been identified as the causative agent of skin lesions in horses and goats, and exudative epidermitis (greasy pig disease) in swine (15, 16, 17, 18). In addition, *S. hyicus* isolates from exudative epidermitis of pigs were found to produce epidermolytic exfoliative toxins, which damaged the superficial layer of the skin (19, 20).

The foregoing observations led to the hypothesis that bacterial infections could also have a role in the pathogenesis of BF lesions. In this study, we isolated and identified *Staphylococcus spp*. from BFs and BF lesions from different north Australian beef herds, sequenced the genomes of selected *Staphylococcus* isolates, and investigated the presence of virulence factors in various isolates to assess the potential role of bacteria in the development of BF lesions.

## MATERIALS AND METHODS

### Sample collection

Samples were collected from cattle with ulcerated to partially scabbed BF lesions from four different herds. Lesions around the eye, on the shoulder, dewlap, and belly of cattle, were swabbed using Amies agar gel transport swabs (Copan, Murrieta, CA, USA). Multiple lesion swabs were collected if the animal had more than one lesion. The cattle sampled included five Brahman heifers from herd 1 (H1) and six Brangus steers from herd 2 (H2), kept at the University of Queensland Pinjarra Hills Research Precinct (−27.53, 152.91) in spatially separated paddocks sampled in May 2020. Eight Brahman heifers from a commercial cattle property in north Queensland (H3) (−20.50, 146.0) sampled in August 2020, and eight Droughtmaster cattle kept at Darbalara farm (−27.59, 152.38) (H4) sampled in January 2021. One swab was also collected from visually normal skin from each animal (at least 15 cm away from lesions). Lesion swabs from five heifers from H1 and two steers from H2 were sampled again in August and September 2021, respectively, and skin swabs were also collected from 15 cm down the medial canthus from three steers without any lesions from H2. This study was approved by the University of Queensland Animal Ethics committee (Approval no. 2021/AE000054).

Buffalo flies were also collected from the back of each swabbed animal with lesions from H1, H2 and H4 and the three steers without lesions from H2 using an insect collection net. The insect net was disinfected with 80% ethanol between each collection and BFs from each collection were transferred to a sterile plastic zip bag, labelled with the animal ID, and transported to the laboratory on ice.

### Bacterial isolation

Each swab was streaked onto 5% (v/v) sheep blood agar and MacConkey agar within 24 hours of collection. Buffalo flies were streaked following the method described previously (21). Briefly, five BFs from each animal were washed three times by dipping into 250 μl of sterile normal saline, and the saline washing solution from each group was streaked onto blood agar and MacConkey agar. After washing, the flies were surface disinfected by immersion in 10% sodium hypochlorite (NaClO) followed by 70% ethanol for at least 10 minutes each. Flies were then rinsed in normal saline, air-dried, and homogenized in 100 μl of sterile normal saline and plated on blood agar and MacConkey agar plates. Plates were incubated at 37°C for 24 hours. Bacterial colonies were distinguished as Gram-positive or negative by the potassium hydroxide (KOH) test (22).

### Species identification by MALDI-TOF

Pure cultures of all isolates from the year 2020 sampling were submitted to the Biosecurity Queensland Veterinary Laboratories (Department of Agriculture and Fisheries, Coopers Plains 4108, Queensland Australia, −27.55, 153.04) for initial species identification by matrix-assisted laser desorption ionization-time of flight mass spectrometry (MALDI-TOF MS) (Bruker Daltonics, Germany).

### DNA extraction

DNA was extracted using the Qiagen DNeasy^®^ Blood and Tissue extraction kit (QIAGEN Pty Ltd., Hilden, Germany) according to the manufacturer’s protocol. Briefly, extraction involved suspending a loop full of the bacterial isolate from a fresh overnight blood agar culture into lysis buffer. The suspension was incubated for 1.5 hours at 56°C. The remainder of the protocol was as recommended by the manufacturer. DNA concentration was measured using a NanoDrop spectrophotometer (NanoDrop 2000, Thermofisher Scientific, MA, USA).

### Genotyping by rep-PCR

A repetitive sequence-based PCR (rep-PCR) was performed as described by Versalovic et al., using primers REP 1R-IDt and REP 2-IDt (REP_1R-IDt: 5′ NNNNCGNCGNCATCNGGC 3′ and REP_2-IDt: 5′ NCGNCTTATCNGGCCTAC 3′) (23). Briefly, the PCR was performed in a total volume of 25 μl containing 5 X GoTaq^®^ buffer (Promega, Madison, WI, USA), 2.5 mM MgCl2, 6.25 mM of each dNTP, 50 pM of each primer 2U of GoTaq^®^ DNA polymerase (Promega, Madison, WI, USA) and 100 ng of DNA template. The cycling conditions included initial denaturation at 95°C for 7 min, annealing at 42°C for 60 sec and *Taq* polymerase activation at 65°C for 8 min followed by 33 cycles of denaturation, annealing and extension at 94°C for 60 s, 42°C for 30 s, and 65°C for 8 m for the final extension, respectively. The reaction was conducted in an Eppendorf Mastercycler^®^ pro thermal cycler (Eppendorf AG 22331 Hamburg, Germany). The PCR products were run on the 2%-TAE buffer (40mM tris, 20mM acetate and 1mM EDTA, pH 8.5) agarose gel containing 0.05 μg/ml of ethidium bromide (Sigma Aldrich, Saint Louise, MO, USA), for 3.5 h at 70V.

The rep-PCR profiles of 44 isolates (from 2020) were compared using BioNumerics software (version 4.50, Applied Maths Inc., Saint-Martens-Latem, Belgium). Genotypic profiles were compared using band matching tolerance and optimization of 0.5%. For cluster analysis of DNA fingerprinting data, the similarities were calculated using the Dice similarity coefficient (24). A comparison dendrogram was also developed using the unpaired group method with arithmetic average (UPGMA). The genotyping of 19 isolates from 2021 were compared with 21 isolates (one representative isolate from each cluster) collected in 2020, using BioNumerics software with the same band matching tolerance and optimization of 0.5%.

### Selection of isolates for sequencing

The 44 *Staphylococcus* isolates from 2020 were grouped into 21 clusters using a cut-off threshold of 90% similarity in rep-PCR based genotyping. A total of 21 isolates were selected (one representative from each cluster) for whole-genome sequencing using the Illumina NovaSeq 6000 (Illumina, San Diego, CA, USA) platform. The selected isolates included three isolates from BFs, two isolates from normal cattle skin and 16 isolates from BF lesions. Nine of the 21 isolates sequenced with the Illumina platform, including eight isolates sourced from lesions and one sourced from BFs, were also sequenced using Oxford Nanopore Technology (ONT) to generate complete genomes. These nine isolates were selected at a cut-off threshold of 70% similarity of the genotypic patterns.

### DNA extraction and quality testing for sequencing

Selected bacterial isolates were sub-cultured from storage (−80°C) by inoculation onto 5% (v/v) sheep blood agar. The cultures were incubated for 24 h at 37 °C. For genome sequencing, DNA was extracted using Gentra Puregene Core Kit A (QIAGEN Pty Ltd., Hilden, Germany) with some modifications of the manufacturer protocol. A standard suspension from the blood agar (OD600 =1.85 or 1.74×10^9^ bacteria) was prepared in sterile phosphate buffer saline (PBS), and an aliquot of 300 μl was transferred to a 2 ml tube. For efficient lysis, 50 μl of lysozyme (Sigma-Aldrich, St. Louis, MO, USA) at a final concentration of 2.9 mg/ml and 50 μl of lysostaphin at a final concentration of 0.14 mg/ml (Sigma-Aldrich, St. Louis, MO, USA) were also added to each tube and incubated at 37°C for 3 h in a thermomixer (Eppendorf Thermomixer C, Hamburg, Germany). After 3 h incubation, 500 μl of cell lysis solution was added to each tube and incubated for 1 h at 56°C. To maximize the DNA yields, the tubes were incubated for 5 m at 80°C. The remaining protocol was followed as recommended by the manufacturer. DNA for sequencing was quantified with Qubit™ 4 fluorometer (Thermo Scientific, Waltham, MA, USA) using Qubit™ dsDNA BR assay kit (Invitrogen, Waltham, MA, USA).

Before library preparation, DNA quality was tested by Pippin Pulse Field Gel Electrophoresis. For this, 500 ng of the extracted DNA samples were run on 0.75% Seakem^®^ gold agarose gel (Lonza Bioscience, Basel, Switzerland) for 16 hours on Pippin Pulse Electrophoresis Power Supply System (Sage Science, Beverly, MA, USA) at waveform type 5-430 kb 80 V. The gel was stained with the 1X concentration of SYBR Safe DNA Gel Stain (Invitrogen, Waltham, MA, USA).

### Illumina library preparation and sequencing

DNA extracted from 21 isolates were sequenced at the Australian Centre for Ecogenomics (ACE), The University of Queensland (St. Lucia, 4072, Queensland, Australia). Briefly, DNA libraries were prepared according to the manufacturer’s protocol using the Nextera DNA Flex Library Preparation Kit (Illumina, San Diego, CA, USA). Library preparation and bead clean-up was undertaken with the Mantis^®^ Liquid Handler (Formulatrix, Bedford, MA, USA) and Epmotion (Eppendorf, Hamburg, Germany) automated platform. These programs cover “Tagment Genomic DNA” to “Amplify DNA” in the protocol (Mantis-Nextera DNA Flex library prep protocol, Illumina, San Diego, CA, USA) and “Clean Up Libraries” in the protocol (Epmotion - Library Clean Up protocol, Eppendorf, Hamburg, Germany). On completion, each library was quantified, and quality control performed using the Quant-iT™ dsDNA HS Assay Kit (Invitrogen, Waltham, MA, USA) and Agilent D1000 HS tapes (Agilent Technologies, Santa Clara, CA, USA) on the TapeStation 4200 (Agilent Technologies, Santa Clara, CA, USA), as per the manufacturer’s protocol.

Nextera DNA Flex libraries were pooled at equimolar amounts of 2 nM per library to create a sequencing pool. The library pool was quantified in triplicate using the Qubit™ dsDNA HS Assay Kit (Invitrogen, Waltham, MA, USA). Library quality control was performed using the Agilent D1000 HS tapes on the TapeStation 4200 as per the manufacturer’s protocol. The library was prepared for sequencing on the NovaSeq 6000 (Illumina, San Diego, CA, USA) using NovaSeq 6000 SP kit v1.5, 2 x 150 bp paired-end chemistry, according to the manufacturer’s protocol.

### ONT library preparation

For ONT sequencing, a MinION sequencing library was prepared using the Nanopore Ligation Sequencing Kit (Oxford Nanopore Technologies, Oxford, UK) as per the manufacturer’s protocol with starting DNA amount of 6 μg (185 fmol). A final amount of 650 ng (20 fmol) of the prepared library was loaded on MinION Spot on Flow Cell (Oxford Nanopore Technologies, Oxford, UK, versions FLO-MIN106D R9.4.1) using flow cell priming kit (Oxford Nanopore Technologies, Oxford, UK) as instructed by the manufacturer. The library was sequenced using a MinION device (Mk1C, MC110367) for 5-6 h with default instrument settings. Primary acquisition of data and real-time base calling was carried out using the graphical user interface MinKNOW (version 20.10.6, Oxford Nanopore Technologies, Oxford, UK) and Guppy basecaller (v4.5.2, Oxford Nanopore Technologies, Oxford, UK).

### Sequence analysis, assembly, and annotation

All data analysis in this study was performed using programs on Galaxy Australia (https://usegalaxy.org.au/). For each isolate sequenced with the Illumina 6000 NovaSeq platform, 1.0 Gbps of data (400X coverage) was acquired in FASTQ format. Initial read quality was determined by FastQC (Galaxy version, 0.72 + Galaxy1) (http://www.bioinformatics.babraham.ac.uk/projects/fastqc). Low-quality reads were removed using Trimmomatic (Galaxy version 0.36.6) (25) and trimmed from the start (LEADING) and at the end (TRAILING) of the reads if the quality score fell below 30. A sliding window trimming was done if the average quality of four bases dropped below 20, and all unpaired reads and reads below 30 bps were removed. Paired reads were *de novo* assembled using the Shovil assembler (Galaxy version 1.1.0+galaxy0) (with settings as; “Estimated genome size”:2.5 Mbps, “Minimum contig length”: 500, “Assembler”; Spades) (https://github.com/tseemann/shovil).

For the isolates sequenced in duplicate with ONT, 2.0 Gbps data (800X coverage) was acquired in FASTQ format. Reads were concatenated using the tool “Concatenate (cat) tail to head” (version 0.1.0 + Galaxy) (https://github.com/bgruening/galaxytools). Reads were filtered for length and average Q score by FiltLong (Galaxy version 0.20 + galaxy1) (https://github.com/rrwick/Filtlong). Reads shorter than 5,000 to 15,000 were removed, depending upon the initial read length N50 histogram (basecalled bases) report generated by Mk1C for an individual isolate. For removal of low-quality reads, a threshold of Q-score ≥ 7 (for 100% reads) and Q-score ≥ 12 (for more than 80% reads) was used. The filtered ONT reads along with respective corrected Illumina reads of the same isolate were used to generate hybrid *de novo* assemblies with Unicycler (Galaxy version 0.4.8.0) with normal “Bridging mode” and “Pilon” option enabled for assembly polishing (26). The *de novo* assembled genome sequences were annotated with the prokaryotic genome annotation tool “Prokka” (Galaxy Version 1.14.6+galaxy0) (27).

### Pangenome analysis, reads mapping, and multilocus sequence phylogeny

To identify the extent of genomic diversity in the sequenced isolates, a pangenome analysis was performed using “Roary” (Galaxy Version 3.13.0+galaxy1) (28). Output from Roary was uploaded on an online web-based platform for interactive visualization of genome phylogenies “Phandango” (29) to visualize the presence and absence of a gene and genomic similarity between isolates.

To confirm the identification of the sequenced isolates, corrected paired Illumina reads from all the sequenced isolates were mapped against genomes of reference strains of *S. agnetis* (DSM23656), *S. hyicus* (NCTC10350), *S. chromogenes* (NCTC10530) and *S. sciuri* (NCTC12103) using Bowtie2 (Galaxy Version 2.4.2+galaxy0) (30). Accession numbers and the strain type for the genome sequences used for reads mapping are listed in Table 1.

**Table 1:** A list of all the bacterial sequences and their corresponding accession numbers used in genome mapping and phylogenetic analysis

| Species name | Type strain | Accession number |
| --- | --- | --- |
| *Staphylococcus agnetis* | DSM23656 | PPQF01000001 |
| *Staphylococcus agnetis* | 1379 | CP045927 |
| *Staphylococcus hyicus* | NCTC10350 | LS483304 |
| *Staphylococcus chromogenes* | NCTC10530 | UHDB01000002 |
| *Staphylococcus sciuri* | NCTC12103 | LS483305 |
| *Staphylococcus aureus* | ATCC25923 | CP009361 |
| *Staphylococcus argenteus* | DSM28299 | JADAMU010000003 |
| *Staphylococcus caprae* | NCTC12196 | UHCW01000006 |
| *Staphylococcus epidermidis* | ATCC14990 | CP035288 |
| *Staphylococcus intermedius* | NCTC11048 | UHDP01000003 |
| *Bacillus subtilis* | NCIB3610 | CP020102 |

A multilocus sequence phylogeny was constructed using nucleotide sequences of four housekeeping genes (*tuf, rpoA, rpoB, recN*) from all the annotated genomes from this study and each of the reference strains of *S. agnetis, S. hyicus, S. chromogenes, S. sciuri, S. aureus, S. caprae, S. epidermidis, S. intermedius, S. argenteus* and *Bacillus subtilis*. The nucleotide sequences for these four housekeeping genes were concatenated (in the order *tuf, rpoA, rpoB, recN*) and aligned for each genome using Geneious software (version 2021.1.1, Biomatters Ltd., Auckland, New Zealand). Accession numbers and the strain type for the genome sequences used for phylogenetic analysis are listed in Table 1.

Before phylogenetic analysis, a model test was performed in Mega X (31) with the aligned nucleotide sequences of the above-mentioned genes to select the best suitable model for phylogeny. The model with the lowest BIC scores (Bayesian Information Criterion) was selected for further analysis. A phylogenetic tree based on concatenated sequences of *tuf, rpoA, rpoB* and *recN* was inferred using the Maximum Likelihood method and General Time Reversible model (32) using Mega X (31). Initial tree(s) for the heuristic search were obtained automatically by applying the Neighbor-Join and BioNJ algorithms to a matrix of pairwise distances estimated using the Maximum Composite Likelihood (MCL) approach. The topology with the superior log likelihood value was then selected. A bootstrap consensus tree inferred from 1000 replicates was taken to represent the evolutionary history of the taxa analyzed (33). Branches corresponding to partitions reproduced in less than 70% bootstrap replicates were collapsed. This analysis involved 32 nucleotide sequences. All positions containing gaps and missing data were eliminated (complete deletion option). There was a total of 7,359 nucleotide positions in the final dataset.

### Virulence factor (VF) identification

To determine how the isolated bacteria contribute to the development of skin lesions, a comprehensive VF search was undertaken to identify potential VF resembling previously reported VF among Staphylococcal and non-Staphylococcal species, including those previously shown to cause skin infections.

To identify the VF sequences, present in each bacterial isolate, a comprehensive virulence factor (VF) data set of staphylococci (CVFS) developed by Naushad et al., (34) was used. Briefly, this custom database was created from the amino acid sequences of the VF for genus *Staphylococcus* obtained from publicly available databases including the Victor database (http://www.phidias.us/victors/), the PATRIC database (35), the VFDB database (36) and phenol-soluble modulins sequences from the UniProtKB database (37). A cut-off of ≥ 30% amino acid sequence similarity, ≥ 50% query length coverage, and ≥ 100-bit score was used to confirm VF presence or absence in the genomes of the corresponding isolates (38, 39, 40).

For robust identification of VF and to identify any VF resembling *non-Staphylococcus* species in our isolates, we assigned identity based on a cut-off ≥ 50% amino acid sequence similarity, ≥ 50% query length coverage, and ≥ 100-bit score, with the amino acid sequences of VF core dataset (VFCD) from VFDB (36). This was conducted using custom BLAST databases for CVFS and VFCD formatted using “NCBI BLAST + makeblastdb” (Galaxy Version 2.10.1+galaxy0) (42, 43, 44) and homology of all amino acids identified in the annotated genomes determined using “NCBI BLAST + BLASTp” (Galaxy Version 2.10.1+galaxy0) (41, 42, 43). As two databases were used for the designation of gene names, to prevent any duplication of gene annotation, the amino acid query sequence with the highest similarity score obtained was used for further analysis.

### Exfoliative toxin analysis

The relationships occurring between the exfoliative toxins identified from isolates in this study and other publically available amino acid sequences for exfoliative toxins A (*eta*) and C (*etc*) of *Staphylococcus* spp. was determined by amino acid sequence alignment in Geneious (version 2021.1.1, Biomatters Ltd., Auckland, New Zealand) and phylogenetic analysis. For *eta*, amino acid sequences for exfoliative toxin A from *S. hyicus* (accession no. AB036768), and *S. aureus* strains MSSA476, N315, NCTC8325, USA300, MRSA252 and RF122 (VFCD id; VFG004880, VFG004849, VFG004878, VFG004876, VFG004881 and VFG004877, respectively) were aligned with *eta* sequences from the current study in Geneious (version 2021.1.1, Biomatters Ltd., Auckland, New Zealand). For *etc*, amino acid sequences from *S. aureus* (accession no. BAA99412), *S. sciuri* (accession no. JF755400), and *S. hyicus* (accession nos. AF515455 and KT072730) were aligned with *etc* sequences identified from this study using the same software.

To select the best suitable model for phylogenetic analysis of *eta* and *etc* genes, a model test was performed in Mega X (31) with amino acid sequences from both genes, and the model with the lowest BIC scores (Bayesian Information Criterion) was chosen for further analysis. Phylogenetic trees were inferred for *eta* and *etc*, respectively, using the Maximum Likelihood method and General Reversible Chloroplast model (44) in Mega X. Initial tree(s) for the heuristic search were obtained automatically by applying the Neighbor-Join and BioNJ algorithms to a matrix of pairwise distances estimated using the JTT model. The topology with a superior log likelihood value was then selected. The bootstrap consensus tree inferred from 1000 replicates was taken to represent the evolutionary history of the taxa analyzed (33). Branches corresponding to partitions reproduced in less than 70% of the bootstrap replicates were collapsed. For the *eta* and *etc* proteins, 27 and 25 amino acid sequences and 306 and 280 amino acid positions were involved, respectively.

### PCR based identification of isolates

To identify the isolates from 2021 sampling and to confirm the identity of isolates from 2020 which were not utilised for genome sequencing, DNA samples extracted from each of the isolates were tested with an *aroD* based multiplex PCR using the primers reported by Adkins et al., for species-specific identification of *S. hyicus* (aroD_hyF: 5′ TATGGTGTCGACCAATCGAAGGCT 3′ and aroD_hyR:5′ ACCCTATAGCCCGCTTAC-TT 3′) and *S. agnetis* (aroD_agF: 5′ CGCATGAGAGACCAATACGCT 3′ and aroD_agR:5′ TAGGACGTATAGAGGTGG 3′) (45). Briefly, the PCR reaction was performed in a total volume of 20 μl containing 10 μl of 2X Phusion Hot Start II High-Fidelity PCR Master Mix (Thermo Scientific, Waltham, MA, USA), 10 μM of each forward and reverse primers and 3 ng of DNA template. The cycling conditions included an initial denaturation at 98°C for 30 s, followed by 30 cycles of denaturation, annealing and extension at 98°C for 10 s, 60°C for 30 s and 72°C for 30 s, respectively, and a final extension at 72°C for 10 m. The reaction was set up in an Eppendorf Mastercycler^®^ pro thermal cycler. For visualisation, the amplification products were run on 2% TBE-buffer (89 mM Tris, 89 mM boric acid, 2 mM EDTA pH 8) agarose gel for 75 m at 80V using GeneRuler 100 bp plus DNA ladder (Thermofisher Scientific, Waltham, MA, USA).

## RESULTS

### Bacterial isolation and identification by MALDI-TOF

Lesion swabs produced small, round, white, non-hemolytic, Gram-positive staphylococcal-like colonies on blood agar. Swabs from active lesions yielded pure cultures, while swabs from partially active lesions produced mixed cultures with dominant growth of staphylococcal-like colonies.

All the six lesion swabs from H1 produced growth of an organism identified as *S. hyicus* by MALDI-TOF, with four swabs having a pure culture. Swabs from normal skin of the H1 heifers did not produce any bacterial growth resembling that seen with the lesion swabs. Buffalo fly surface rinses from one animal yielded three colonies of *S. hyicus*, while pure cultures of *S. hyicus* were isolated from homogenized BFs plated from two animals with lesions. All eight lesion swabs from H2 also yielded *S. hyicus*, with pure cultures obtained from six lesions. Three swabs from normal skin of H2 steers (including one from a non-lesion steer) produced one to two colonies of *S. hyicus* with abundant environmental contaminants, whereas one swab from the normal skin of an animal with lesions yielded two colonies of *S. sciuri*. A pure culture of *S. hyicus* was isolated from homogenized BFs collected from two steers (one with lesions and one without lesions). Twelve swabs from H3 produced *S. hyicus* colonies, with very heavy growth from four lesion swabs. Staphylococcal-like colonies of varying sizes were present on seven swabs from H3. Multiple colonies (one representative from each size variant) were purified by sub-culture. Eight lesion swabs collected from H4 also yielded staphylococcal growth in blood agar, and the bacteria were isolated as pure cultures from three swabs. A normal skin swab from one H4 animal produced only one colony of *Staphylococcus* spp., while BFs washings from one animal and homogenized BFs from two animals yielded two staphylococcal-like colonies and heavy staphylococcal-like growth, respectively. All swabs collected from H1 and H2 in 2021 had staphylococcal-like growth, with pure cultures grown from two swabs. The MALDI-TOF technique was not used to identify any isolates from the 2021 sampling from herds H1, H2 and H4. All the 2020 and 2021 isolates were re-identified by PCR, and no bacterial growth was observed following plating on MacConkey agar.

### Strain typing by rep-PCR

Strain typing by rep-PCR was completed on all 44 isolates collected in 2020 from lesions, BFs and unaffected skin which had been identified by MALDI-TOF as described above. From these 44 isolates, 21 different banding patterns (cluster/pattern type 1-21) were identified at a 90% similarity cut-off value. Overall, ten clusters contained a single isolate, five clusters contained two isolates, and the remaining six clusters contained more than two isolates. Thirty-five isolates from the lesions belonged to 17 different clusters, of which six clusters consisted of a single isolate. Five isolates from the BFs were placed in three clusters, with three isolates being in a cluster with a lesion isolate, while two isolates belonged to single isolate clusters. Four isolates from normal skin showed four different strain types, one of which was *S. sciuri* as distinguished based on a banding pattern different from most of the other isolates. Only one isolate from the normal skin had a similar strain pattern, with one lesion isolate from the same animal.

Among the 19 isolates collected in 2021, 15 different patterns were observed (cluster/pattern type 22-36), of which 13 were identified only once. Upon comparison, none of the 2021 isolates showed strain similarity with any of the 21 strain types isolated in 2020. The same pattern was observed from normal skin on one animal, one BFs, and two lesion isolates from 2021 collections. There were no differences between the genotypic patterns of isolates from different herds. The details for the cluster/pattern type of each isolate are provided in Table 2.

**Table 2:**
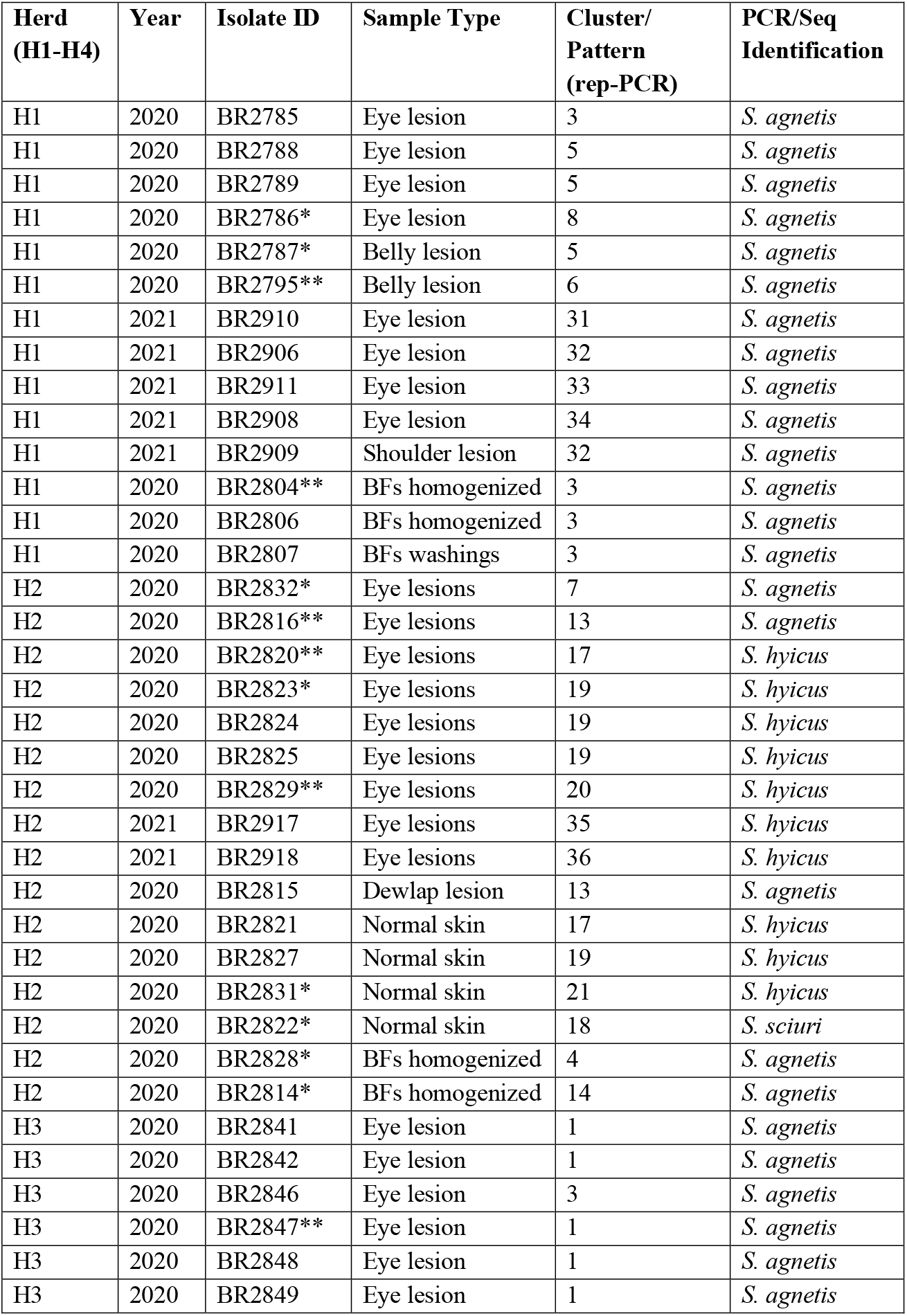
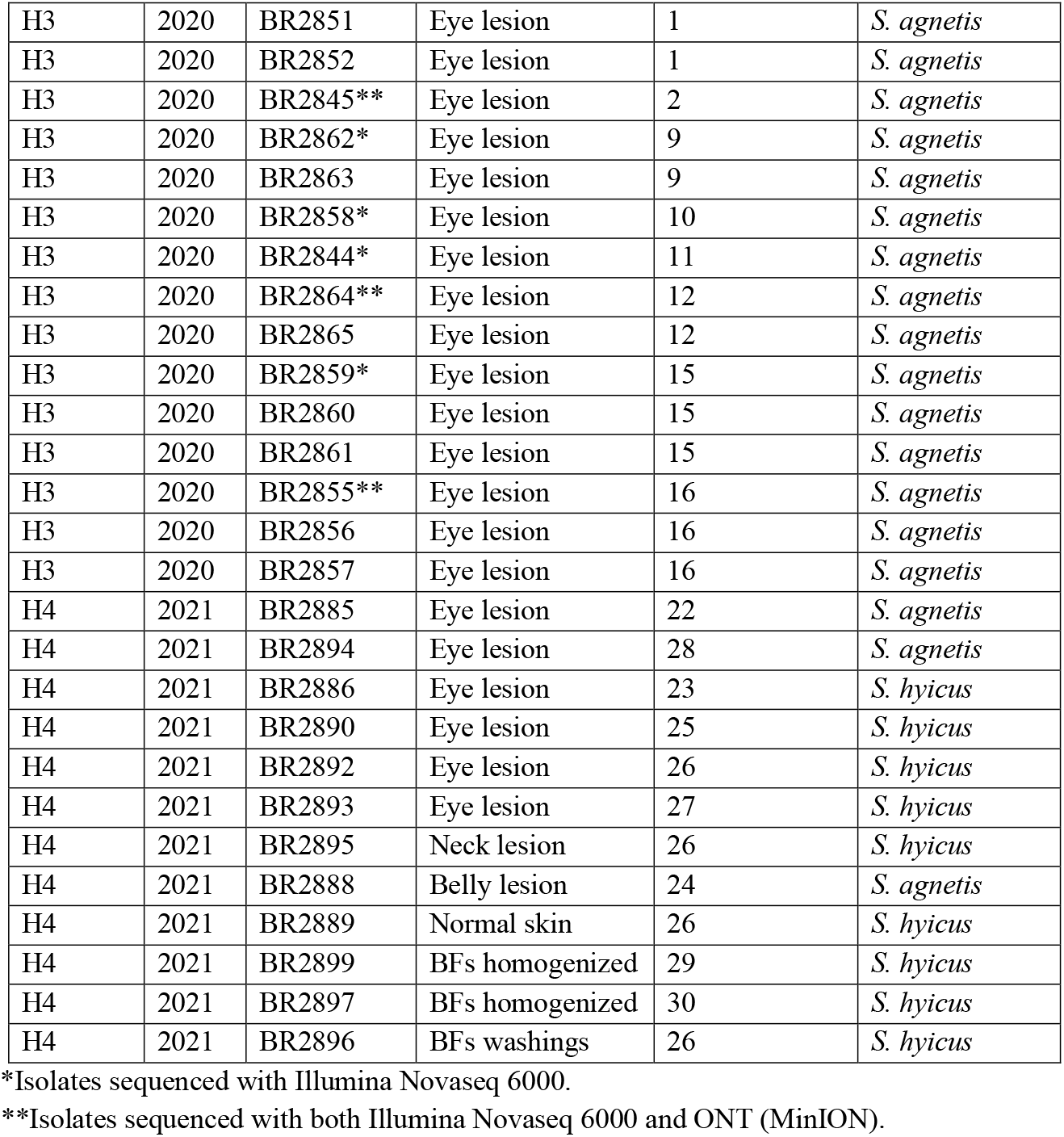
Details of isolation, source, cluster/pattern type and PCR/Seq based identification of bacterial isolates from the current study.

### Genome assemblies and annotation

To genetically characterize and determine the role of the bacterial isolates associated with lesion development, 21 isolates (one representative from each cluster/pattern type) from the 2020 sampling were selected for whole-genome sequencing and virulence factor analysis (Table 2). In this study, we generated draft, *de novo* genome assemblies for 12 representative isolates (source: Lesion = 8, BFs = 2, Normal skin = 2) from 2 x 150 bp paired Illumina reads. For the rest of the 9 representative isolates (source: Lesion = 8, BFs = 1), we generated complete finished, *de novo* genome assemblies by hybrid assembly approaches using Illumina and ONT reads. The details of assembly status, number of contigs, genome size, number of CDS, rRNA and tRNA for each isolate sequenced are provided in Table 3. The draft genome assemblies ranged from 38 to 115 contigs, comprising 2.40 to 2.78 Mbps. The hybrid assemblies comprised a single circular contig and range from 2.41 to 2.50 Mbps in size. The number of coding sequences (CDS), ribosomal RNAs (rRNA), and transfer RNAs (tRNA) varied between isolates and ranged from 2334-2769, 3-19 and 42-75, respectively (Table 3). The sequence data generated and analysed in the current study were deposited in the NCBI Genome database under BioProject Accession number PRJNA809943.

**Table 3:** Details of bacterial genome assembly, number of contigs, genome size, number of CDS, rRNA and tRNA for each isolate sequenced in this study. The suffix _aL, _aB, _sN, _hL and _hN after each isolate ID indicate *S. agnetis* from lesion, *S. agnetis* from buffalo flies, *S. sciuri* from normal skin, *S. hyicus* from lesion and *S. hyicus* from normal skin, respectively.

| Isolate ID | Assembly status | Genome size (bp) | No. of contigs | Longest contig (bp) | CDS | rRNA | tRNA |
| --- | --- | --- | --- | --- | --- | --- | --- |
| BR2786_aL | Draft | 2481539 | 89 | 304262 | 2402 | 5 | 48 |
| BR2787_aL | Draft | 2482413 | 112 | 304101 | 2403 | 7 | 58 |
| BR2795_aL | Complete | 2449123 | 1 | NA | 2465 | 19 | 59 |
| BR2832_aL | Draft | 2533020 | 90 | 304049 | 2470 | 6 | 55 |
| BR2844_aL | Draft | 2440290 | 60 | 225370 | 2358 | 6 | 56 |
| BR2845_aL | Complete | 2437387 | 1 | NA | 2371 | 19 | 60 |
| BR2847_aL | Complete | 2414453 | 1 | NA | 2423 | 19 | 59 |
| BR2855_aL | Complete | 2481102 | 1 | NA | 2465 | 19 | 60 |
| BR2858_aL | Draft | 2404324 | 53 | 318300 | 2334 | 3 | 42 |
| BR2859_aL | Draft | 2472985 | 69 | 194377 | 2402 | 3 | 43 |
| BR2862_aL | Draft | 2434216 | 76 | 229021 | 2359 | 6 | 46 |
| BR2864_aL | Complete | 2438730 | 1 | NA | 2373 | 3 | 42 |
| BR2816_aL | Complete | 2506912 | 1 | NA | 2517 | 19 | 59 |
| BR2804_aB | Complete | 2462691 | 1 | NA | 2539 | 19 | 60 |
| BR2814_aB | Draft | 2492807 | 88 | 303910 | 2429 | 4 | 51 |
| BR2828_aB | Draft | 2485945 | 88 | 181960 | 2426 | 4 | 42 |
| BR2820_hL | Complete | 2434473 | 1 | NA | 2390 | 19 | 59 |
| BR2823_hL | Draft | 2597767 | 38 | 489936 | 2556 | 4 | 72 |
| BR2829_hL | Complete | 2452563 | 1 | NA | 2369 | 19 | 62 |
| BR2831_hN | Draft | 2519724 | 76 | 270422 | 2448 | 4 | 75 |
| BR2822_sN | Draft | 2784109 | 135 | 251211 | 2760 | 3 | 43 |

### Pangenome analysis, read mapping, and multilocus sequence analysis (MLSA)

The initial pangenome analysis of all 20 sequenced *S. hyicus* isolates, indicated that there were 754 core genes (99% <= strains <= 100%), 3 softcore genes (95% <= strains < 99%), 3429 shell genes (15% <= strains < 95%), and 2024 cloud genes (0% <= strains < 15%). The smaller number of core genes and the high number of cloud genes indicates that the 20 sequences of isolates initially identified as *S. hyicus*, might actually be different *Staphylococcus* species, rather than just strains within a species. A pangenome analysis for all sequenced isolates is presented in Fig 1.

**Fig. 1:**
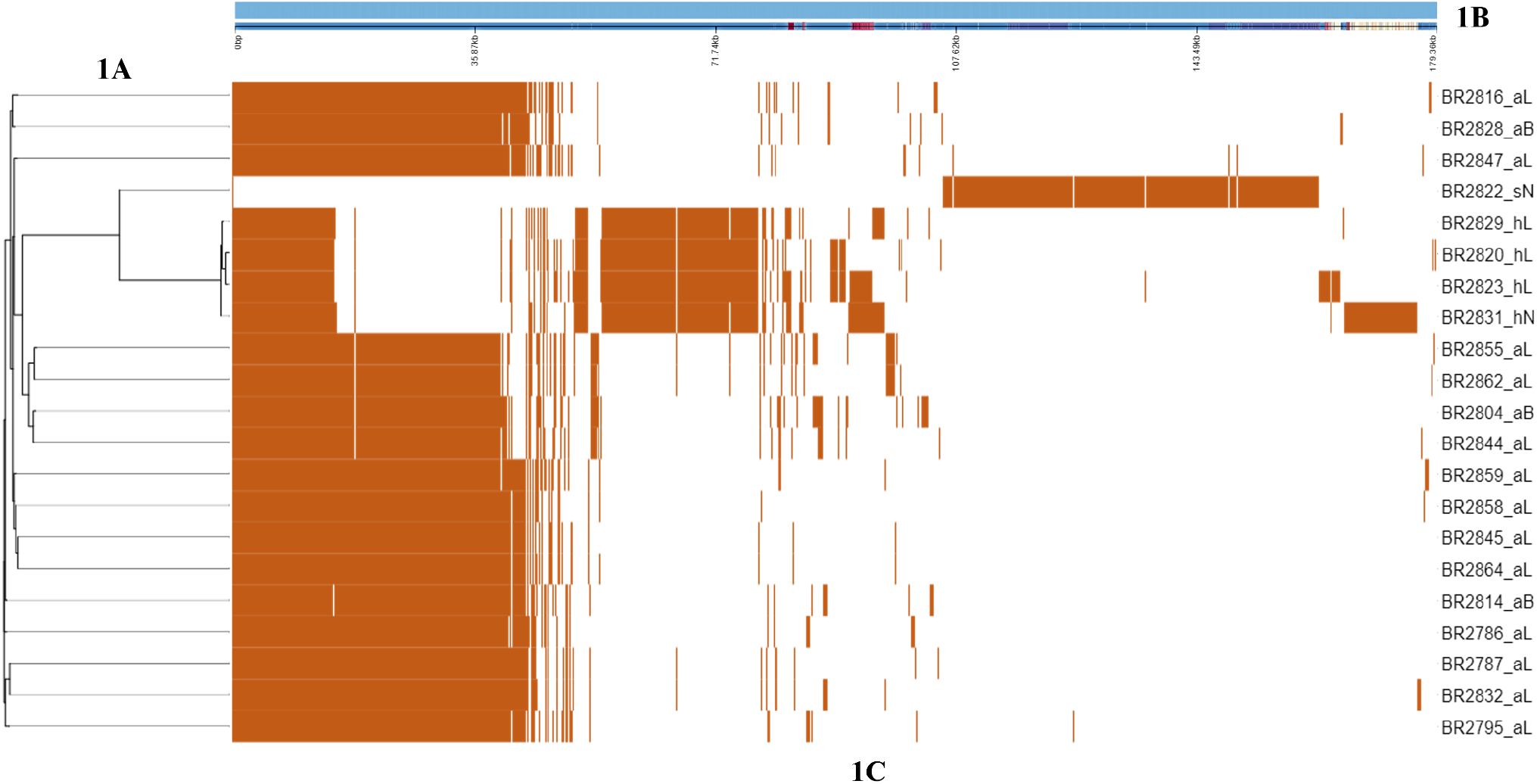
The pangenome analysis indicates a similar and unique gene group among all the sequenced isolates with a core genome phylogenetic tree (1A). The topmost panel (1B) shows a single representative nucleotide sequence inferred for each gene of the pangenome. The central panel (1C) shows the presence (burnt orange), or absence (white) of blocks relative to the genes and contigs in the pan-genome. The suffix _aL, _aB, _sN, _hL and _hN after each isolate ID indicate *S. agnetis* from lesion, *S. agnetis* from buffalo flies, *S. sciuri* from normal skin, *S. hyicus* from lesion and *S. hyicus* from normal skin, respectively.

To confirm the sequence isolate identity, corrected paired Illumina reads from each isolate sequenced were mapped against the genomes of reference strains of *S. hyicus, S. agnetis, S. chromogenes* and *S. sciuri*. Of the 21 sequenced isolates, 80.5% to 90.32% of paired Illumina reads from 16 isolates were mapped with *S. agnetis*, while <30% and <10% of the reads from these 16 isolates were mapped with reference genomes of *S. hyicus* and *S. chromogenes*, respectively. From the remaining isolates, 78-82% of the reads from four isolates mapped with *S. hyicus*, but <30% and <10% of the reads from these four isolates mapped with reference genomes of *S. agnetis* and *S. chromogenes*, respectively. The 89.5% reads from the *S. sciuri* isolate obtained from this study, mapped exactly with the reference genome of *S. sciuri*.

The argument for reclassifying 16 of the *Staphylococcus* isolates was further strengthened when a pangenome analysis of these 16 suspected *S. agnetis* isolates resulted in 1981 core genes, zero softcore gene, 806 shell genes and 896 cloud genes. This finding indicated that these 16 isolates were *S. agnetis*. Pangenome analysis of the four confirmed *S. hyicus* isolates, resulted in the identification of 810 core genes, zero softcore genes, 4211 shell genes and zero cloud genes, indicating more strain variation among these isolates.

The identity of 21 sequenced isolates was confirmed by a multilocus sequence phylogenetic analysis based on four housekeeping genes (tuf, rpoA, rpoB and recN) (Fig. 2). This showed that all the 16 suspected *S. agnetis* isolates clustered (97% branch threshold of homology) with the reference strain of *S. agnetis* (DSM23656) and bovine mastitis isolate of *S. agnetis* (1379). This confirmed that these 16 isolates from the current study belonged to the *S. agnetis* species. Four additional *S. hyicus* isolates clustered strongly (99% branch threshold of homology) with the reference strain of *S. hyicus* (NCTC10350), while one *S. sciuri* isolate from the current study clustered with the reference strain of *S. sciuri* (NCTC12103) (Fig. 2).

**Fig. 2:**
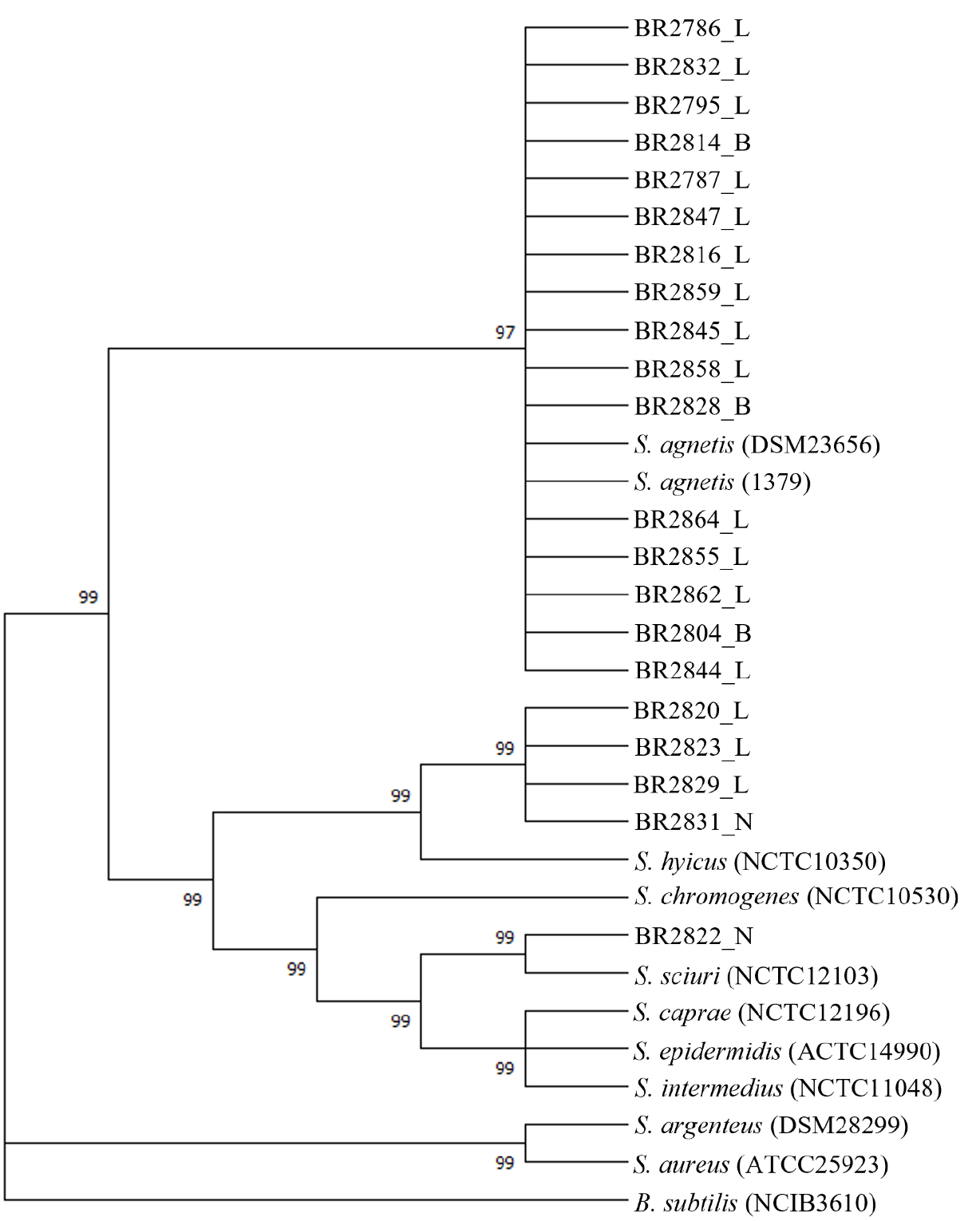
Maximum likelihood multilocus tree built from concatenated nucleotide sequence alignment of four housekeeping genes (*tuf, rpoA, rpoB* and *recN*) of all the isolates and other closely related *Staphylococcus* spp. Bootstrap branch support (based on likelihood analysis) is shown. The suffix _L, _B and _N, after each isolate ID indicate “from lesion”, “from buffalo flies” and “from normal skin”, respectively.

### Staphylococcal VF identification

We identified 13 genes that belonged to eight VF of the adherence category, including autolysin (*atl*), clumping factors (*clfA* and *clfB*), collagen adhesion (*cna*), fibrinogen binding proteins (*efb*), fibronectin-binding proteins (*fnbA* and *fnbB*), intracellular adhesin (*icaA, icaB* and *icaC*), Ser-Asp-rich fibrinogen binding proteins (*sdrD* and *sdrF*) and staphylococcal protein A (*spa*). The distribution and percentage similarities for the staphylococcal VF genes identified are provided in Fig 3. The distribution of the VF varied between the different species isolated as well as between different strains within species. The autolysin gene (*atl*) was the only gene identified in all the *Staphylococcus* isolates for which genome sequencing was undertaken. The VF genes responsible for adherence identified in the genome sequences of all the *S. agnetis* isolates were almost the same, except that the *clfA, clfB, sdrD* and *sdrF* genes were present in 87.5%, 93.75%, 75% and 62.5% of the isolates, respectively. Similarly, VF genes responsible for adherence identified in the genome sequences of the *S. hyicus* isolates were also the same except that *clfA, efb*, and *sdrD* were absent in the single isolate obtained from normal skin sample. The staphylococcal protein A gene (*spa*) was found only in the *S. hyicus* isolates whereas the genes *clfA, clfB, cna, efb, fnbA, fnbB, sdrD* and *sdrF* were not identified in the *S. sciuri* isolate, which instead contained *icaA, icaB* and *icaC* genes.

**Fig. 3:**
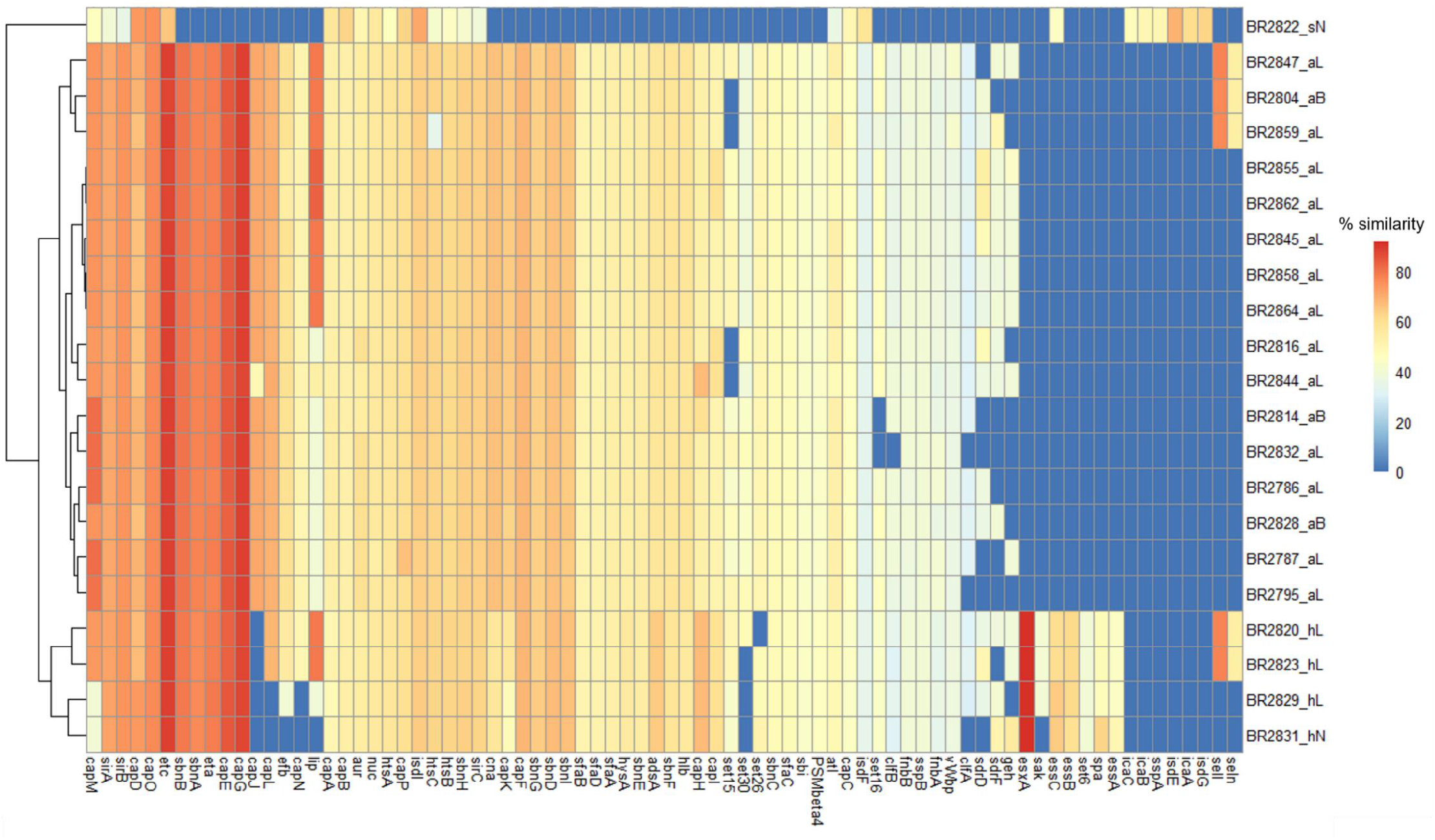
Heat map showing distribution and amino acid sequence similarities of different virulence factors (VF) genes in the 21 sequenced isolates from 2020, with the VF sequences of *Staphylococcus* spp. in the databases. The dendrogram indicates clustering of isolates based on the presence or absence and percentage similarities of different VF. The genes for adherence category include *atl, clfA, clfB, can, efb, fnbA, fnbB, icaA, icaB, icaC, sdrD, sdrF*, and *spa*. The genes *aur, adsA, sspB, geh, lip, sspA, sak, nuc*, and *nuc* are belonged to exoenzymes type VF. The genes responsible for host immune evasion include capsule forming genes (*capA, capB, capC, capD, capE, capF, capG, capH, capI, capJ, capK, capL, capM, capN, capO* and *capP*) and staphylococcal binder of immunoglobulin (*sbi*). The genes responsible for iron uptake and metabolism include *isdE, isdF, isdG, isdI, htsA, htsB, htsC, sfaA, sfaB, sfaC, sfaD, sirA, sirB, sirC, sbnA, sbnB, sbnC, sbnD, sbnE, sbnF, sbnG, sbnH* and *sbnI*. The genes involved in type VII secretion system include *essA, essB, essC* and *essD*. The gene *seln* and *sell* are encoded for exotoxins while *set6, set15, set16, set26, set30* are for enterotoxin. The exfoliative toxin A and C are encoded by the gene *eta* and *etc* respectively. The gene *hlb* and *PSMβ4* are hemolysin and phenol soluble modulins genes respectively. The suffix _aL, _aB, _sN, _hL and _hN after each isolate ID indicate “*S. agnetis* from lesion”, “*S. agnetis* from buffalo flies”, “*S. sciuri* from normal skin”, “*S. hyicus* from lesion” and “*S. hyicus* from normal skin”, respectively.

Among the exoenzymes examined, we identified nine VF including aureolysin (*aur*), adenosine synthase A (*adsA*), cysteine protease (*sspB*), lipases (*geh* and *lip*), serine V8 protease (*sspA*), Staphylokinase (*sak*), thermonuclease (*nuc*) and von-Willebrand factor binding protein (*vWbp*). Of these, *aur* and *nuc* were the only genes identified in all the sequenced *Staphylococcus i*solates. Most of the exoenzyme genes identified within the *S. agnetis* isolate were similar, except for the gene *geh* found in 68.75% of the isolates. Similarly, VF genes for exoenzymes were mainly similar in the *S. hyicus* isolates, except for the genes *geh, lip* and *sak*, which were present in 75%, 75% and 50% of the isolates, respectively. The gene *sspA* was present in only the *S. sciuri* isolate.

The VF category involved in host immune evasion consists of genes for capsule formation (*capA, capB, capC, capD, capE, capF, capG, capH, capI, capJ, capK, capL, capM, capN, capO* and *capP*) and the staphylococcal binder of immunoglobulin (*sbi*). The gene *sbi* was identified in the genome sequences of all isolates, except that of the *S. sciuri* isolate. All the capsule forming genes classified within the host immune evasion VF category, were identified in the genome sequence of all the *S. agnetis* isolates characterized. The gene *capJ* however, was absent in all *S. hyicus* isolates, while *capL* and *capN* were not found in 50% of the genome sequences for this species. In addition, the *S. sciuri* isolate had only *capA, capB, capC, capD, capM, capO* and *capP* genes for capsule formation within the whole genome sequence.

For iron uptake and metabolism VF categories, 23 genes were identified including four iron-regulated surface determinant genes (*isdE, isdF, isdG* and *isdI*), seven ATP-binding cassette (ABC) transporter (siderophore receptors) genes (*htsA, htsB, htsC, sfaA, sfaB, sfaC and sfaD*), three staphyloferrin A synthesis related genes (*sirA, sirB* and *sirC*) and nine staphyloferrin B synthesis related genes (*sbnA, sbnB, sbnC, sbnD, sbnE, sbnF, sbnG, sbnH* and *sbnI*) (Fig. 3). All of the genes responsible for iron uptake and metabolism were identified in all *S. agnetis* and *S. hyicus* isolates except *isdE* and *isdG*, which were only identified in *S. sciuri*. The genes *sbnA, sbnB, sbnC, sbnD, sbnE, sbnF, sbnG, sbnI, sfaA, sfaB, sfaC* and *sfaD* were not identified in *S. sciuri*.

The four genes for the type-VII secretion system (*essA, essB, essC* and *essD*) were identified in all *S. hyicus* isolates, while *S. sciuri* had only one gene (*essC*). No type VII secretion system gene was found in any of the *S. agnetis* isolates. The beta-hemolysin gene (*hlb*) was identified in all *S. agnetis* and *S. hyicus* isolates but absent in *S. sciuri*. The genes for enterotoxin (*sell* and *seln*) were identified in 50% of *S. hyicus*, and 18.75% *S. agnetis* isolates, but absent in *S. sciuri*. All the isolates of *S. agnetis* and *S. hyicus* were found to have exfoliative toxin A and C genes (*eta* and *etc*), while the *S. sciuri* isolate had only the *etc* gene. The genes for exotoxin (*set26* and *set30*) were identified in all the *S. agnetis* isolates, while the genes *set15* and *set16* were only present in 81.25% and 87.5% of the isolates, respectively. All *S. hyicus* isolates carried *set6, set15, set16* genes for exotoxins, while the genes *set26* and *set30* were found in 75% and 25% of the isolates. The phenol soluble modulins gene (*PSMβ4*) was identified in all *S. agnetis* and *S. hyicus* isolate but no exotoxin or phenol soluble modulins genes were observed in *S. sciuri*.

### Non-staphylococcal VF identification

We also identified some VF in our isolates that had ≥ 50% amino acid homology with VF of non-staphylococcal species in VFCD (Fig. 4). Among these, 22 genes of the VF enzyme category were detected including urease (*ureA, ureB* and *ureG*), 6-phosphogluconate dehydrogenase (*gnd*), catalase (*katA*), adenylylsulfate kinase (*cysC1*), ATP-dependent Clp protease proteolytic subunit (*clpP*), capsule biosynthesis protein (*capC*), chaperonin (*groEL*), endopeptidase Clp ATP-binding chain (*clpC*), flagellar-related 3-oxoacyl-ACP reductase (*flmH*), glutamate-1-semialdehyde-2,1-aminomutase (*hemL*), molecular chaperone (*ct396*), autolysin (*aut*), nitrate reductase (*narH*), nucleoside diphosphate kinase (*ndk*), pantoate--beta-alanine ligase (*panC*), UTP--glucose-1-phosphate uridylyltransferase (*bpsC*), undecaprenyl diphosphate synthase (*uppS*), phosphopyruvate hydratase (*eno*), prolipoprotein diacylglyceryl transferase (*lgt*) and protein disaggregation chaperone (*clpB*). All isolates had almost all the above-mentioned VF genes except the *aut* gene, which was absent in three *S. agnetis*, all *S. hyicus* and *S. sciuri* isolates. The gene cysC1 was also absent in all *S. hyicus* and *S. sciuri* isolates. All isolates had three additional genes for iron uptake and metabolism, including *fagC, vctC, cpsJ* whereas the 4^th^ gene *iraT* was only found in three *S. hyicus* isolates from lesions. Three genes for putative proteins (*plr/gapA, hpt* and *lplA*) were identified in all *S. agnetis* isolates, while *S. hyicus* and *S. sciuri* exhibited only *plr/gapA* and *lplA*.

**Fig. 4:**
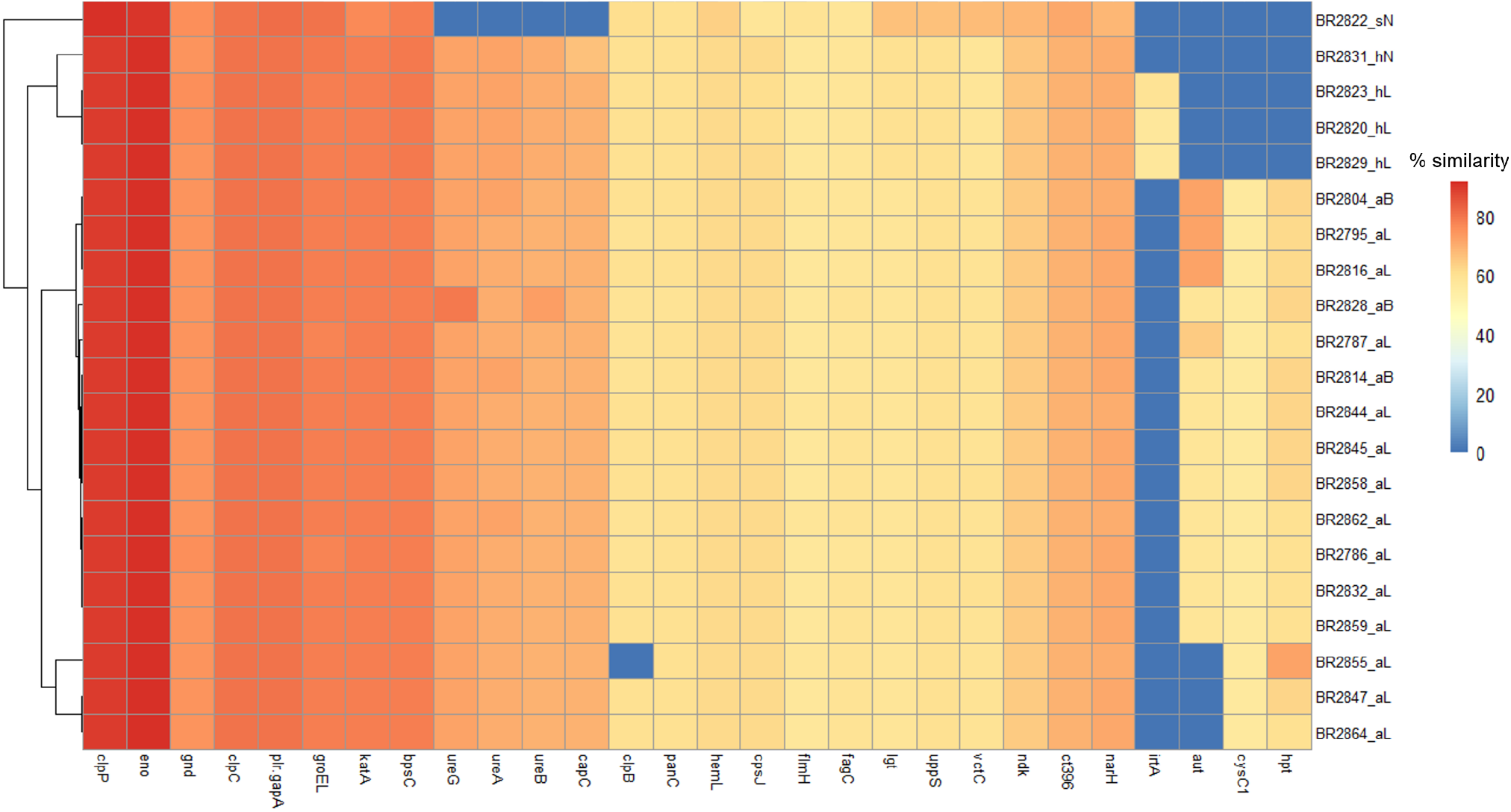
Heat map showing distribution and amino acid sequence similarities of different virulence factor (VF) genes in all sequenced isolates with the VF sequences of non-staphylococcal species in the databases. The dendrogram indicates clustering of isolates based on the presence or absence and percentage similarities of different VF. This includes genes for enzyme VF category (*ureA, ureB, ureG, gnd, katA, cysC1, clpP, capC, groEL, clpC, flmH, hemL, aut, narH, ndk, panC, bpsC, uppS, eno*, and *lgt*) additional genes for iron uptake and metabolism (*fagC, vctC, cpsJ, iraT*). The genes *plr/gapA, hpt* and *lplA* are encoded for putative proteins. The suffix _aL, _aB, _sN, _hL and _hN after each isolate ID indicate *“S. agnetis* from lesion”, “*S. agnetis* from buffalo flies”, *“S. sciuri* from normal skin”, “*S. hyicus* from lesion” and “*S. hyicus* from normal skin”, respectively.

### Exfoliative toxin analysis

Exfoliative toxin A (*eta*) identified from all *S. agnetis* isolates had 89.21%-89.54% and 77.77%-78.10% amino acid homology with *ExhA* from *S. hyicus* and *eta* from different strains of *S. aureus*, respectively. The *eta* gene from all *S. hyicus* isolates had 94.12%-95.08% and 78.75%-79.73% homology with *ExhA* from *S. hyicus* and *eta* from different strains of *S. aureus*, respectively. The *eta* gene was not identified in the *S. sciuri* isolate from normal skin.

Evolutionary tree analysis for the *eta* gene showed all the *eta* sequences from the current study occurred in the same clade as *S. hyicus ExhA*, but on different branches (Fig 5A). Exfoliative toxin C (*etc*) identified from all the isolates had 87.09%-87.61% amino acid homology with *etc* from *S. aureus*. The *etc* gene from all the isolates had <1% similarity with *S. hyicus* and *S. sciuri*. Evolutionary tree analysis of *etc* gene showed all the *etc* sequences from the current study occurred in the same clade as *S. aureus*, but on different branches (Fig. 5B).

**Fig. 5:**
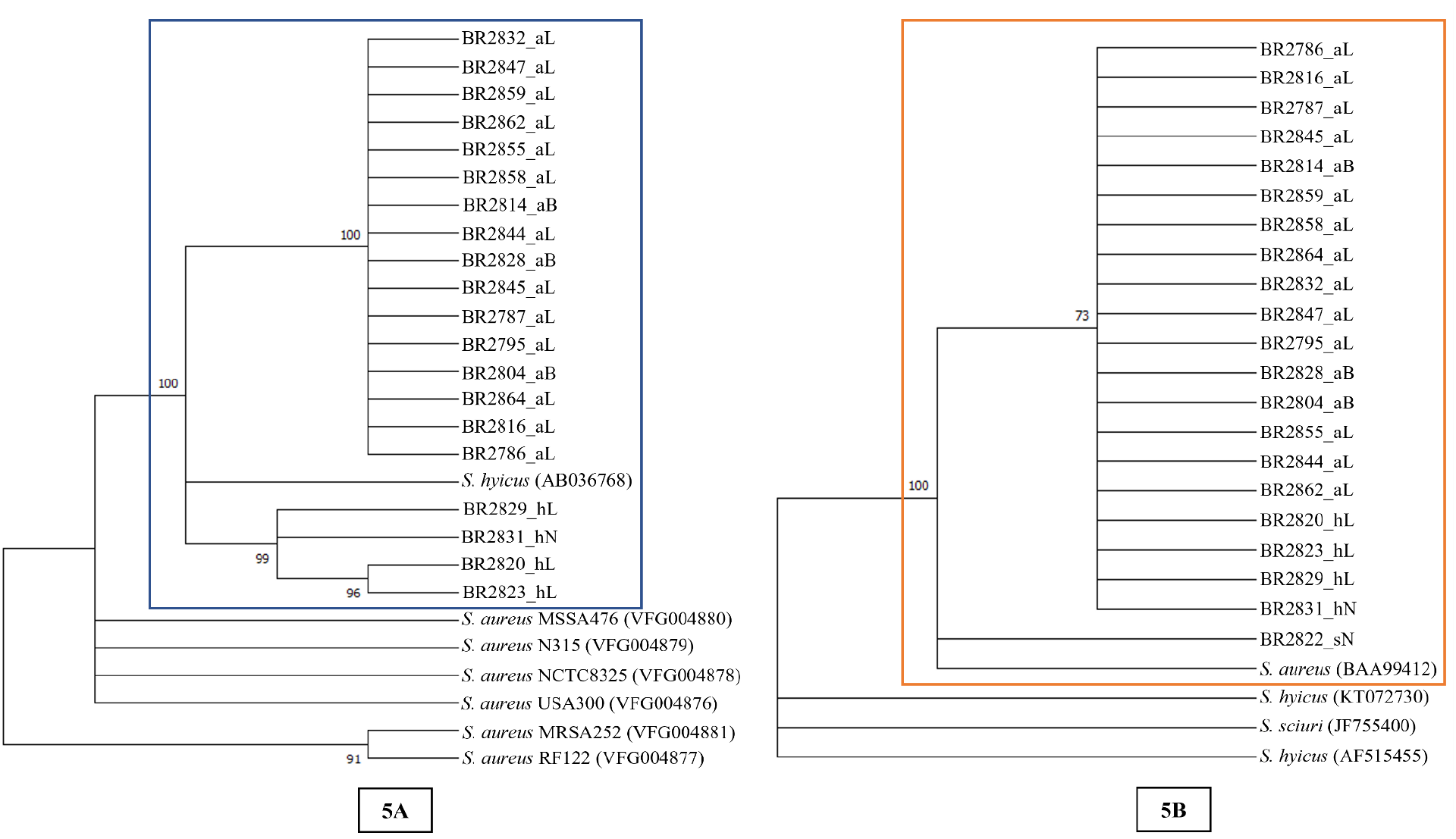
Maximum likelihood trees built from amino acid sequences of *eta* (5A) and *etc* gene (5B) from all the sequenced isolates and other closely related staphylococcal species. Bootstrap branch support (based on likelihood analysis) is shown. The suffix _aL, _aB, _sN, _hL and _hN after each isolate ID indicate *“S. agnetis* from lesion”, *“S. agnetis* from buffalo flies”, *“S. sciuri* from normal skin”, “*S. hyicus* from lesion” and “*S. hyicus* from normal skin”, respectively.

### PCR based identification

As MALDI-TOF was unable to differentiate *S. agnetis* from *S. hyicus*, an *aroD* gene-based species-specific PCR amplifying 295 bps for *S. agnetis* and 425 bps for *S*. *hyicus* was used for re-identification of all isolates. In initial identification by MALDI_TOF, all 62 isolates were identified as *S. hyicus*, and one normal skin isolate was identified as *S. sciuri*. But re-identification by *aroD* gene-based PCR identified 43/62 isolates as *S. agnetis* and only 19/62 as *S. hyicus*. All the lesion and BFs isolates collected from H1 in 2020 and 2021 were identified as *S. agnetis*. From H2, seven lesions and three normal skin isolates were identified as *S. hyicus*, while three lesions and two BFs isolates were identified as *S. agnetis*. Two lesion isolates from the year 2021 were also identified as *S. hyicus*. All 20 lesion isolates from H3 were confirmed as *S. agnetis* with species-specific PCR. From H4, five lesion isolates, three BFs isolates (two from homogenized BFs and one from washings) and one normal skin isolate were identified as *S. hyicus*, whereas three lesion isolates were confirmed as *S. agnetis*. The details of isolation, source and identification of bacterial isolates are provided in Table 2.

## DISCUSSION

In the past, the development of *Haematobia-associated* lesions has generally been attributed to the effects of infection with *Stephanofilaria* spp. nematodes transmitted by BF and HF, but in a number of studies, the failure to find nematodes in all lesions has suggested that other causal factors, including bacteria, may also be involved (9, 11). In this study we identified *S. agnetis* and *S. hyicus* from BFs and BF lesions from different north Australian beef herds using whole genome sequencing and conducted a subsequent comprehensive VF identification, which indicated a potential role of these bacterial species in lesion pathogenesis.

All BF lesions sampled in this study were found to be infected with either *S. agnetis* or *S. hyicus*, and the bacteria were isolated in pure cultures from fresh lesions. *Staphylococcus hyicus* has previously been reported as a causative agent of skin lesions and teat infection in cattle (13, 11, 46), and exudative epidermitis in pigs (18). Devriese and Derycke, suggested that *S. hyicus* skin infection in cattle occurred secondary to parasitic infestation (13) but Hazarika et al., reproduced the skin lesions in rabbits by experimental inoculation of *S. hyicus* isolated from cattle skin lesions (14). Al-Rubaye et al., indicated the potential of *S. agnetis* to cause skin lesions following the identification of exfoliative toxin genes similar to *S. hyicus* isolates of swine-origin and *S. aureus* of the scaled-skin syndrome in humans (47). Both *S. agnetis* and *S. hyicus* have been isolated as causative agents of bovine mastitis and intra-mammary infections (11, 45, 48).

We isolated *S. agnetis* in pure culture from surface-sterilised homogenised BFs from four cattle, and a single colony of *S. agnetis* was also isolated from BFs exocuticle washings from one animal. Similarly, *S. hyicus* were isolated in pure cultures from BFs from two animals with lesions whereas one BFs washing sample yielded two colonies of *S. hyicus*. Horn flies have also been reported to vector *S. aureus* and transmit bacteria into the teat skin, resulting in the development of abscesses and lesions (8, 10). Anderson et al., isolated *S. aureus* from 55.8% of the HF collected from three herds with *S. aureus* intramammary infection (49) and Owens et al., found that *S. aureus* can remain active in infected HF without a significant change in the bacterial count for up to four days (8). In a 16S rDNA-based pyrosequencing microbiome study of the HF, Palavesam et al., identified *S. hyicus* in adult male HFs and HF eggs (12). This 2012 study (12) did not identify any *S. agnetis* in HFs, although this could be because *S. agnetis* was first described as separate species to *S. hyicus* in 2012 (48).

The 43 *S. agnetis* isolates in our study included 23 different strain types and the 19 *S. hyicus* isolates were of 12 different strain types. Four strains of *S. agnetis* were isolated from multiple animals and one of these strains was isolated from lesion samples from two separate herds. Two strains of *S. hyicus* were also identified from BF lesions from multiple animals. Hazarika et al., collected 47 *S. hyicus* isolates from cattle skin lesions which were separated into 10 different strains by biochemical characterization (14). Wegener et al., also reported that the pigs with exudative epidermitis were infected with multiple strains of *S. hyicus* (18). Similarly, 42 *S. agnetis* isolates from a mastitis study of a dairy herd showed 23 different banding patterns with pulsed-field gel electrophoresis (PFGE) strain typing (45). Isolation of multiple strains of *S. agnetis* and *S. hyicus* from BF lesions in combination with previous reports (14, 18, 45) of multiple strain involvement of these species in skin and mammary gland infections, suggests that there might be multiple strains of these two species involved in the pathogenesis of BF lesions.

The genotypic pattern of 75% *S. agnetis* isolates (three of 4 isolates) from BFs collected from animals with lesions were identical to lesion isolates from two herds, whereas out of three *S. hyicus* isolates from BFs, only one (33.33%) showed similarity with two lesion isolates. In a study of mastitis in three herds, Anderson et al., identified eight different genotypic patterns from 244 isolates of *S. aureus* from teats, milk/colostrum and horn flies (49). Of the *S. aureus* isolates from HF, 82.7% belonged to a single genotypic group and 51.6% had an identical genotypic pattern to the mastitis isolates. Similarly, Gillespie et al., noted that the *S. aureus* isolates from HFs had similar genotypic patterns to the 95% of *S. aureus* isolates from udder/teat infections in heifers (11), whereas in our study, 50% of BFs isolates had identical genotypic patterns to the BF lesion isolates. None of the *S. hyicus* isolates from the normal skin had a similar genotypic pattern to the lesion isolates from the respective animals except for one instance, and no *S. agnetis* were isolated from the normal skin samples in our study. The isolation of pure cultures of *S. agnetis* and *S. hyicus* from the surface sterilised homogenised BFs that were similar to lesion isolates suggests that BFs might play an important role in the transmission of these bacteria.

The inability of MALDI-TOF to differentiate between *S. hyicus* and *S. agnetis* species noted in our study has also been reported previously (50, 51). In our study, this was due to the lack of *S. agnetis* in the MALDI-TOF database used, which meant that a lower percentage of isolates were correctly identified despite a MALDI-TOF score of ≥2.0 when this species was not in the database. MALDI-TOF is based on the analysis of 16S rDNA gene sequences, which might not be useful to differentiate these two species as 16S rDNA gene sequences of *S. agnetis* isolates from the current study showed 99.87-99.92% similarity with *S. hyicus* NCTC10350, which is higher than the recommended cut-off value of 98.7% similarity for different bacterial species (52). Taponen et al., also reported 99.7% similarity of 16S rDNA gene sequences between *S. agnetis* isolates and *S. hyicus* ATCC11249 (49). The complete sequencing of the β subunit of RNA polymerase (*rpoB*) gene and the elongation factor Tu (*tuf*) gene have been used previously to differentiate *S. hyicus* and *S. agnetis* at significantly higher similarity cut-off values (≥97% and ≥98%, respectively) (45, 53). However, from a cross-species gene similarity comparison we identified that the DNA repair protein gene (*recN*) of *S. agnetis* isolates has 99.34-99.76% and 82.32-82.44% similarity with *S. agnetis* DSM23656 and *S. hyicus* NCTC10350, respectively, while *recN* gene of *S. hyicus* isolates has 98.75-99.10% and 82.14-82.44% similarity with *S. hyicus* NCTC10350 and *S. agnetis* DSM23656, respectively. These apparent differences in the *recN* gene sequence similarities of these two species indicate that the *recN* gene can also be used as a potential marker to differentiate these two species.

The 78 different virulence factor genes identified in sequenced isolates in this study were known virulence factors from the genus *Staphylococcus*, and 28 genes had been identified in bacterial species other than *Staphylococcus*. The genes for adherence are almost similar in both *S. agnetis* and *S. hyicus* isolates except for staphylococcal protein A (*spa*) which was only present in *S. hyicus* in this study. This finding was consistent with Naushad et al. (2019) who reported *spa* genes in all three bovine mastitis *S. hyicus* isolates but not in 13 *S. agnetis* isolates (34). The presence of clumping factor B gene (*clfB*) in all our *S. agnetis* and *S. hyicus* isolates was the only difference between the adherence genes identified in this study and those from the bovine mastitis study of Naushad et al., where only 15% of bovine mastitis *S. agnetis* isolates had this gene (34). The intracellular adhesion genes or biofilm-producing genes (*icaA, icaB, icaC*) were only identified in the *S. sciuri* isolate in our study, which is also consistent with the results of previous studies (34,54).

After adherence to the host surface, bacterial pathogens produce different enzymes which help neutralize the host immune response and promote tissue degradation (55). Our study identified 10 different exoenzymes genes potentially involved in host immune system neutralization, and most of them were common to *S. hyicus* and *S. agnetis* isolated in the current study. The exoenzyme gene profile of isolates from our study was very similar to those previously reported from bovine mastitis isolates of these species (34, 56), except that we identified the gene for lipase enzyme (*geh*) in 11 *S. agnetis* and three *S. hyicus* isolates. Our study also identified three genes responsible for urease activity (*ureA, ureB, ureG*) in all isolates except *S. sciuri*, and this is in line with the study of Åvall-Jääskeläinen et al., (56).

Pathogenic bacteria also use encapsulation to evade the host immune system, and staphylococci are well equipped with encapsulation genes enabling their protection against phagocytosis and enhancing persistence of infection (57, 58). The *S. agnetis* isolates from our study had all of the previously identified encapsulation genes (*capA*-*capP*) except that the *capJ* gene was absent in all isolates and *capL* and *capN* were absent in 50% of the *S. hyicus* isolates. Similarly, Naushad et al., reported the absence of *capN* gene in all isolates of *S. agnetis* and *S. hyicus*, and *capJ* in *S. hyicus* isolates (34). In addition, Åvall-Jääskeläinen et al., did not find *capH-capK* genes in *S. agnetis* isolates from bovine mastitis in Finland (56).

Bacterial pathogens require iron for replication and to maintain infection for which pathogenic bacteria have various iron acquisition mechanisms (59, 60, 61). The profile of iron uptake genes for *S. agentis* and *S. hyicus* isolates from our study was similar to the previous reports (34, 56) with four type-VII secretion system (T7SS) genes in all *S. hyicus* isolates, but none of the *S. agnetis* isolates had these genes, which was the only virulence factor difference we observed between these two species. The T7SS genes encode for a protein secretion pathway, considered important for the virulence of Gram-positive bacteria (62, 63). This finding is similar to that of Åvall-Jääskeläinen et al., who also found lack of T7SS genes in their *S. agnetis* isolates (56). In contrast, Naushad et al., reported six different T7SS genes in 31% and 67% of the *S. agnetis* and *S. hyicus* isolates, respectively (34). Our study also identified the beta hemolysin gene (*hlb*) in all sequenced isolates except *S. sciuri*, which supports the findings of Åvall-Jääskeläinen et al. (56) and Naushad et al. (34).

Identification of exfoliative toxin A (*eta*) and C (*etc*) genes from *S. agnetis* and *S. hyicus* in this study was a major finding in relation to skin lesion development. These toxins are also known as epidermolytic toxins and are serine proteases, able to digest skin desmoglein-1 resulting in the deterioration of desmosomal cell adhesions and epidermal damage (64). The presence of the *eta* gene in *S. agnetis* and *S. hyicus* isolates is in accordance with previous mastitis isolate studies (34, 47, 56), but none of these studies found the *etc* gene in their isolates. Our study also identified the *etc* gene in *S. sciuri* from normal skin, which is contrary to findings of Naushad et al. (34). The close amino acid homology of exfoliative toxins A and C from this study with exfoliative toxins A and C from *S. hyicus* isolated from exudative epidermitis of pigs and *S. aureus* from skin infections in horses, indicates that these toxins might play an important role in the epidermal damage in BF lesions. This suggestion is further strengthened by the observations from histological studies of BF lesions which indicated epidermal damage in all bacterially infected lesions (unpublished observation). Our study also indicated similarity between the VF profiles of BFs and BF lesion isolates, which further suggests that the BFs isolates where the genotype is not similar to the lesion isolate are equally pathogenic and may involve in lesion pathogenesis when transmitted during BF feeding. The close similarity between the VF profiles of *S. agnetis* and *S. hyicus* indicate that both of these species could potentially be involved in the pathogenesis of BF lesions.

## CONCLUSION

The findings from this study indicate that the bacteria *S. agnetis* and *S. hyicus*, vectored by BF, are likely to be significant factors in the pathogenesis of BF lesions. This suggests that the role of bacteria should be a consideration in the development of optimal treatment and control strategies for BF lesions.

